# Resting state brain signal complexity of young healthy adults reflects genetic risk for developing Alzheimer’s Disease

**DOI:** 10.1101/2020.11.08.373167

**Authors:** Xiaojing Li, Yadwinder Kaur, Oliver Wilhelm, Martin Reuter, Christian Montag, Werner Sommer, Changsong Zhou, Andrea Hildebrandt

## Abstract

The e4 allele of the APOE gene is strongly associated with impaired brain functionality and cognitive decline in humans at older age. It is controversial whether and how the APOE e4 allele is affecting brain activity among young healthy individuals and how such effects may contribute to individual differences in cognitive performance. Signal complexity is a critical aspect of brain activity that has been shown to be associated with brain function. In this study, we analyzed multiscale entropy (MSE) of EEG signals among young healthy adults as an indicator of brain signal complexity and investigated how MSE is predicted by APOE genotype groups. Furthermore, by means of structural equation modeling, we investigated whether MSE predicts fluid intelligence. Results indicate larger MSE in young healthy e4 carriers across all time scales. Moreover, better fluid intelligence (gf) is associated with smaller MSE at low time scales and larger MSE at higher scales. However, MSE does not account for better cognitive performance among APOE e4 carriers by mediating the APOE genotype effect on fluid intelligence. The present results shed further light on the neural mechanisms underlying gene-behavior association relevant for Alzheimer’s Disease risk.

## Introduction

Human APOE gene variants are well-known risk factors for cognitive impairment in later life (Michaelson, 2014). The APOE gene has three alleles, e2, e3, and e4, yielding six genotypes (e2/e2, e2/e3, e2/e4, e3/e3, e3/e4, and e4/e4). Abundant evidence suggests that genotype groups including the APOE e4 allele are characterized by reduced brain activity and cognitive decline in the elderly (Reiman et al., 1996; Small et al., 1995; Rubino et al., 2013; Whitehair et al., 2010). However, for young healthy adults, associations of APOE genotypes with cognitive abilities, and brain activity are inconclusive (Li et al., 2019; Weissberger et al., 2018). Available evidence points to a nature-nurture interaction. Among the highly educated population, young carriers of the APOE e4 allele tend to have better cognitive abilities estimated on the basis of multiple tasks as compared with non-carriers, whereas among the people with lower education the e4 allele exhibited lower cognitive abilities than non-carriers (Li et al., 2019). However, whether and how cognitive abilities in young e4 allele carriers relate to brain activity remains to be investigated.

Recently, the concept of entropy, reflecting non-linear spatio-temporal dynamics, has been employed to brain signals. Most studies on young healthy individuals suggest that brain systems with higher signal complexity operate closer to a critical state (He et al., 2013) where a small input may induce a large effect. Thus, it has also been suggested that higher signal complexity indicates a greater dynamic range (Garrett et al., 2013) and improved robustness of the neural system to disruption (Basalyga and Salinas, 2006) and may, therefore, be beneficial for cognition.

In many studies, multiscale entropy (MSE; Costa et al., 2002; 2005) has become a widely applied measure of complexity in neural signals (e.g., Garrett et al., 2013; Takahashi et al., 2010; 2016). MSE is estimated via an advanced algorithm, parameterizing signal complexity at fine to coarse time resolutions, that is, at low to high time scales. At the lowest time scale, MSE extracts information from the entire frequency spectrum of the original signal and thus also captures linear stochastic effects which is dominant in the entropy measure; at higher time scales, MSE captures the slower oscillatory fluctuations of the signal. Thus, the combination of low and high time scales reflects non-linear signal properties. Indeed, associations of MSE and aging-related cognitive performance have been shown to be highly scale-dependent. For example, declined brain functions in healthy elderly were associated with both, reduced MSE at high scales, as well as increased MSE at low scales (McIntosh et al., 2014; Sleimen-Malkoun et al., 2015). Also, a scale-dependent effect pattern was detected among individuals suffering Alzheimer’s Disease (AD) by Mizuno et al. (2010); as compared with healthy controls, MSE of AD patients with impaired cognitive functions was increased at low scales and decreased at high scales.

Hitherto, there is no research addressing the question whether MSE can also differentiate young healthy APOE e4 carriers from non-carriers. Nor are there comprehensive and systematic investigations on how MSE at low to high scales is associated with specific cognitive abilities. For example, an fMRI study by Yang et al. (2014) found no APOE e4 effect on brain complexity, measured during spatial task performance in young adults at multiple spatial regions. However, genotype-related group differences in fMRI-based complexity may be smeared when MSE is integrated across scales when using simple averaging, as customary in previous studies. Furthermore, low time resolution and short time series characterizing many fMRI data may have provided unreliable MSE estimates. As yet, no study has applied MSE analysis to EEG signals in order to assess APOE genotype effects on temporal brain complexity in young healthy adults although the high temporal resolution of EEG signals may provide rich information to be detected by entropy measures across multiple time scales.

In this study, we investigated how APOE genotypes are related with MSE of EEG signals assessed during resting-states. We further studied whether MSE as a neural measure accounts for individual differences in cognitive performance, which was shown to be superior among (highly educated) APOE e4 carriers (Li et al., 2019). In addition, we examined whether the association of the APOE genotype and cognitive performance can be at least partly explained by individual differences in MSE. Since APOE e4 carriers (with higher education) were demonstrated to have better cognitive performance and considering the research findings reviewed above, we expected decreased MSE at lower time scales and increased MSE at higher scales among young e4 carriers relative to non-carriers in our sample in which the participants have relatively high education level. To test these hypotheses, a discovery and a replication sample was used to detect the scalp location of theses associations and to replicate them in an independent sample. Moreover, we related APOE genotypes, MSE and cognitive ability estimates in an integrated statistical model. Going far beyond previous studies, we applied a latent variable approach to MSE (Kaur et al., 2019) as well as to cognitive performance assessments (Raykov, 2006). This psychometric approach is powerful because it integrates MSE across both time scales and spatial regions of interest, and controls for measurement error.

## Method

### Participants and data treatment

#### Participants

Data from two samples, collected in the German cities Greifswald (Sample 1: discovery) and Berlin (Sample 2: replication), obtained with the same procedure for gene analysis and resting state EEG recordings were separately analyzed. Genetic records were available for 63 participants in Sample 1 and 197 participants in Sample 2. Sample 1 served as discovery sample to identify scalp locations of interest (ROI) where the genetic effect on MSE was most strongly present. Because this study is the first one to investigate MSE differences in APOE genotype groups in young adulthood, the scalp location of possible genotype effects was unknown and had to be discovered first. Sample 2 served replication of the genetic effects in the ROIs determined in the discovery sample and was further used to investigate gene-brain-behavior associations. Since the genotype frequency in the two samples was substantially different (50% vs. 26% e4 carriers in Sample 1 and 2, respectively, with Sample 2 being in line with the distribution in the general population), e4 carriers were randomly and repeatedly under-sampled in the data of Sample 1, equating the e4 carrier frequency with the population distributions and in line with Sample 2. The observed e4 frequency differences in Sample 1 might be due to regional differences (Ward et al., 2012; Singh et al., 2006) or to a sampling bias. After resampling, there were 45 individuals in Sample 1 and 197 individuals in Sample 2. Task performance measures were available for 189 participants out of the total sample of 197. Demographic information about the final samples is provided in Table 1.

**Table 1.** Demographic characteristics of the final samples

|  | Sample 1 | Sample 2 |
| --- | --- | --- |
| Mean age (SD) in years | 23.25 (3.35) | 27.63 (5.38) |
| Frequency of female participants | 60% | 51% |
| Frequency of participants with high-school degree | 79% | 74% |
| Frequency of e4 carriers | 26% | 26% |

The genotyping of the APOE alleles was carried out using the same method as described previously (Li et al., 2019). For statistical analysis, each individual was assigned either to the group of e4 carriers (with one or two e4 alleles; dummy coded 1) or non-e4 carriers (dummy coded 0).

#### EEG recordings

For both samples, EEG data were recorded in quiet environments during two 90-s resting state conditions with eyes open and eyes closed, respectively. During recordings with eyes open, participants looked at a fixation cross on the screen. After 90 s, they were presented with a slide that instructed them to close their eyes; after 90 s they were told to re-open their eyes.

For EEG recording in Sample 1, an elastic EEG cap (Easycap, Brain Products, Germany) was mounted with 30 electrodes based on the 10-20 system. One electrode was placed below the left eye to detect vertical eye movements and blinks. The EEG signals were sampled at 250 Hz. For Sample 2, the EEG was recorded with 42 Ag/AgCl electrodes. Of these electrodes, 40 were positioned in the same type of cap according to the 10-20 system (Pivik et al., 1993). Three electrodes, two at the outer canthi and one below the right eye, were used to detect eye movements and blinks. In both samples Impedances were kept < 5 kΩ and the left-mastoid (M1) served as reference. The EEG was amplified with BrainAmp DC amplifiers (Brian Products, Germany) and sampled at 1 kHz. Offline, these signals were down-sampled to 250 Hz.

#### Pre-processing of the EEG data

EEG signal pre-processing was conducted with the same procedure for both samples with the EEGLAB toolbox (v2019.1; Delorme & Makeig, 2004; Widmann et al, 2015). Signals from 24 electrodes that overlapped across the two datasets were submitted to band-pass filtering from 0.1 to 40 Hz. The extended ICA ‘runica’ algorithm (Delorme, Sejnowski & Makeig, 2007) was used to remove eye blink, movement, and electro-cardio artifacts. MARA (EEGLAB plugin; Chaumon, Bishop, & Busch, 2015) was used to reject artefactual components. Then, data were recalculated to average reference with the original reference channel removed.

### MSE calculation

The MSE calculation was based on the algorithm provided by physionet.org, adapted from the original paper proposing the algorithm for calculating MSE (Costa et al., 2002; 2005):

1. At scale ***τ*** ranging from 1-20, obtain a coarse-grained time series 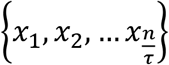 out of the original time series data {*d*_1_, *d*_2_,… *d_n_*}, by calculating the average value of the data points in the window size *τ*, written as Brain signal complexity and APOE polymorphism

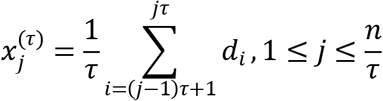 Thus, 20 coarse-grained time series are obtained. For ***τ* =** 1, the time series is the original data.
2. Compute the Sample Entropy (SampEn) of each of the 20 coarse-grained time series, as follows: For a given pattern *i* with length *m* and tolerance *r*, compute the conditional probability *p_i_*(*m*) of the event that all possible series with *m* consecutive data points picked out for the time series that repeats its pattern within distance of *r*, will also repeat for *m* + 1 data points within distance of *r*. MSE is calculated as the negative logarithm of the conditional probability *SampEn*(*m*) = − *lnp*(*m*).

To avoid multiple testing issues in statistical analyses, we integrated the MSE values across adjacent time scales, following the procedure used in a previous study (Kaur et al., 2019), and calculated areas under the curve (AUCs) of MSE as integrated measures at low (1-5), medium (6-11), and high (12-20) time scales, for each participant and electrode. These composite values of MSE were later used to obtain electrode-specific latent variables (see Section 2.4.1).

### Behavioral measures

Raven’s Advanced Progressive Matrices (Rav; Raven and Court, 1979) were used to assess fluid intelligence (gf) in Sample 2. From 16 items of the Rav we created 3 item parcels and estimated a latent gf variable, as previously done to investigate genetic association with gf (Li et al., 2019).

### Statistical analysis

We applied Structural Equation Modeling (SEM) using the R Software for statistical computing (R Development Core Team, 2017) and the lavaan package (Rosseel, 2012). Common standards were used to evaluate the goodness of model fit, with the following values indicating acceptable fit: *χ*^2^ value/df < 2.5, root mean square error of approximation (RMSEA) < .08, comparative fit index (CFI) > .95, and standard root mean square residual (SRMR) < .08.

#### Discovering ROIs for genetic associations of MSE in Sample 1

In order to identify ROIs for later replication analysis, we used the smaller Sample 1 as a discovery sample. For each resting state condition (EO/EC) and each electrode, the area under the curve (AUC) for low (1 – 4), medium (5 – 11) and high (12 – 20) time scales MSE was used to estimate latent MSE variables based on three AUC indicators derived from independent EEG segments. This latent MSE variable was regressed onto the dummy variable coding e4 and non-e4 carriers (Fig. 1A). AUC was calculated according to the procedure proposed by Kaur et al. (2019). The length of segments used for calculating MSE indicators was chosen on the basis of prior reliability analyses (see Supplementary Material S1). As indicated by Figure S1, the reliability of MSE calculated for segments of 22 s duration was satisfactory (reliability > .7) for all scales. Because reliability did not increase much with longer segments, we used 22-s segments to create three MSE indicators for modeling latent variables (see Figure 1A. Regression weights for each latent variable model were standardized to facilitate comparisons across electrodes and time scales.

**Figure 1.**
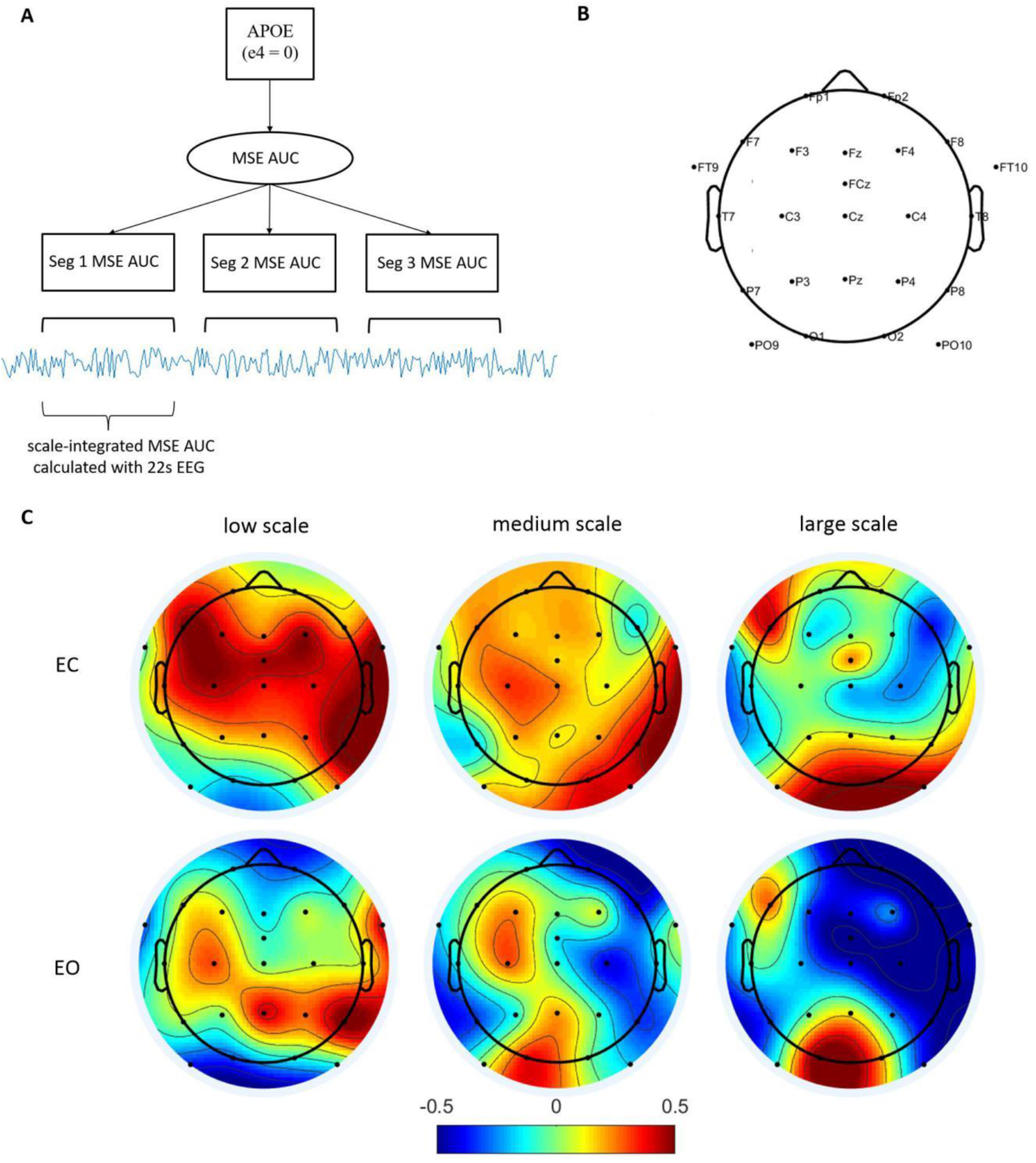
Spatial distributions of the genotype effect for different recording conditions in Sample 1. (A) Schematic representation of the structural equation model exploring APOE genotype associations with MSE at the level of latent variables. Aiming to control for measurement error associated with MSE calculation, the latent variable was obtained from three MSE indicators calculated for different EEG epochs at each electrode and time scale and integrated as AUC across low, medium and high scales. (B) Channel locations for 24 electrodes used for analysis. (C) Topographies of APOE genotype effect sizes on MSE at low, medium and high scale AUC. The red color indicates larger MSE among e4 carriers as compared to non-carriers. EC – eyes closed resting state; EO – eyes open resting state.

#### Replication of the genetic associations of MSE in Sample 2

By using the data from Sample 2, SEM (Model 1) was performed on each resting state condition (EO/EC), three time-scale ranges (AUCs for low, medium, and high scales), and two regions of interest (frontal ROI: F3, F4, Fz; occipito-parietal ROI: O1, O2), as identified in Sample 1. In order to avoid multiple testing issues, we estimated a higher-order latent variable indicated by lower-level electrode-specific latent factors of the electrodes within the ROI. Electrode-specific lower-level factors were indicated by three MSE AUC values, derived from different EEG data segments as explained above in section 2.3.1. The higher-order latent variable was regressed onto the dummy variable, coding e4 carriers vs. non-e4 carriers. A schematic description of the model is shown in Figure 2.

**Figure 2.**
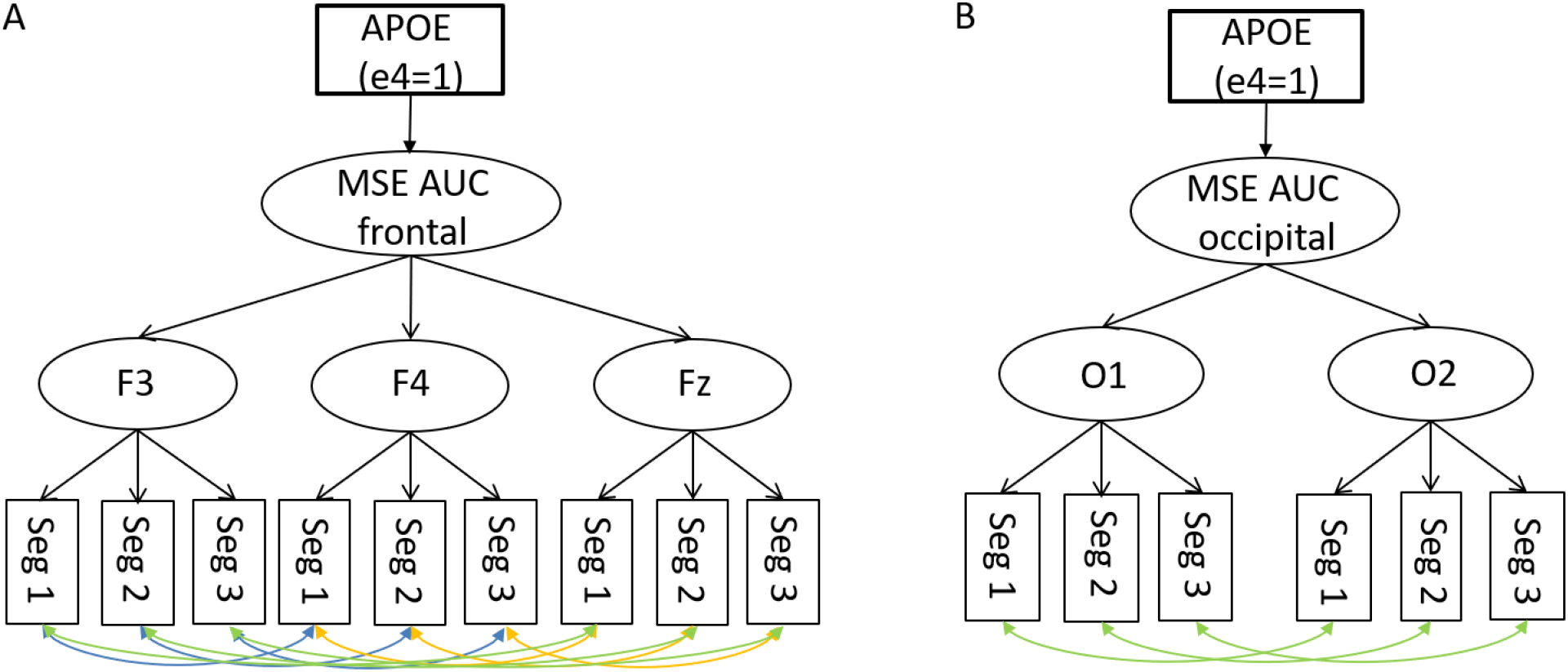
Schematic representations of Model 1 for frontal and occipital ROIs (Panel A and B, respectively) investigating the associations of APOE genotype groups with latent MSE variables for AUC scores and separate EEG data segments (Seg 1 to Seg 3). Indicators are AUC scores calculated from the MSE across the time scales for each segment. Note that these two model versions for the two ROIs were applied to low, medium and high scales in EC vs. EO conditions, resulting in a total of 12 variants of Model 1.

In the models described in Figure 2, there was a high homogeneity among the electrode-specific factors with respect to their association with the higher-order MSE factor. Thus, the loadings of each electrode-specific factor were set to be equal because tau-equivalence was satisfied (Raykov, 1997). Residual correlations were needed to account for similarities still captured by the indicator’s uniqueness if the same time segments (i.e., the first, second vs. third part of the 90-s EEG recordings) were considered among different electrodes. For example, MSE indicators calculated for electrodes F3, F4 and Fz of the frontal ROI in the first EEG segment from start were allowed to correlate in their residuals. The same residual covariances were included for Segment 2 and 3.

#### Investigating the association between MSE and gf in Sample 2

We estimated a further SEM (Model 2) to investigate how gf is associated with MSE. To this purpose, the latent variable gf, indicated by the three Raven item parcels, was regressed onto the latent variable of MSE AUC, as described in 2.4.2, using also the same ROIs. Figure 3A provides a schematic illustration of the model.

**Figure 3.**
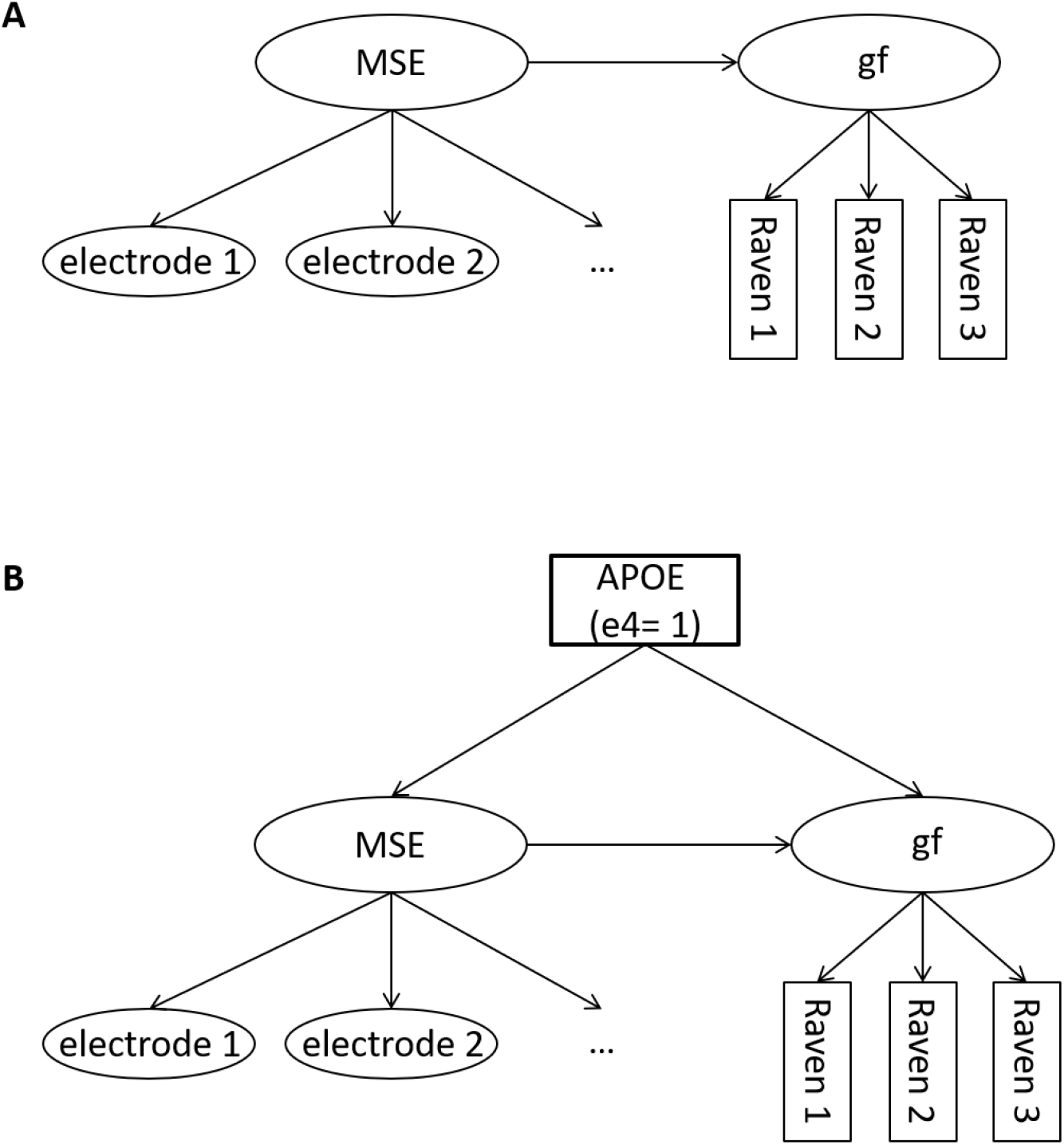
Schematic representation of the structural equation models exploring the MSE effects on gf. Panel A: Model 2, investigating the direct effect of MSE on gf; Panel B: Model 3, investigating the joint relationship between the APOE genotype, MSE and gf, assuming MSE to be a mediator of the gene-behavior association.

#### Does MSE mediate the relationship between APOE and gf ?

Because we expected an association between MSE and gf, we are further interested in whether MSE may mediate the association between APOE and gf which was detected in our previous work (Li et al., 2019). As a mediator, MSE, whether or not independently affecting gf, should diminish or at least weaken the association between APOE and gf. We estimated a third model (Model 3) in which the latent variable of MSE, as described in 2.4.2, was included as a mediator of the regression of gf (modeled as in Model 2) onto the dummy variable coding e4 vs. non-e4 allele carriers. Should the APOE e4 effect on gf diminish after adding MSE as a mediator (Fig. 3B), we can conclude that MSE plays a role in the association between APOE genotype and gf. Figure 3B illustrates Model 3.

## Results

### Identifying ROIs with maximal associations between APOE polymorphisms and MSE

Figure 1C illustrates scalp topographies of effect sizes for e4 vs. non-e4 MSE contrasts at low, medium and high time scale AUCs, separately for the EO and EC conditions, estimated as described in 2.4.1 in Sample 1. The genotype effects in Figure 2A depict regression weights of the standardized latent MSE variable onto the genotype coding dummy variable. In Figure 1C, darker colors indicate larger effect sizes. Positive associations (red color) indicate that APOE e4 carriers show higher signal complexity. During the EC condition, the association was mostly positive across the frontal scalp at low time scales. At medium scales, the spatial pattern of the genetic effect was less systematic, but the effect was more positive at the parietal region. At high scales, there was no genetic effect except at occipital electrodes. In the EO condition, the association was not systematic for low and medium scales, but clearly positive at the occipital region and negative at the centro-frontal region for high scales.

According to the spatial pattern displayed in Figure 1C, we selected the ROIs for further analysis where the e4 effects were largest: electrodes F3, F4 and Fz representing the frontal ROI and electrodes O1 and O2 representing the occipital ROI.

### Replicating the APOE gene association with MSE in an independent sample

After identifying ROIs with associations of APOE polymorphisms and MSE in Sample 1, we aimed to replicate the above association in the selected ROIs in data from Sample 2. We thus estimated latent variables that account for AUC (low scale (time scale 1-5), medium (6-11) and high scale (12 – 20), modeled separately) scores computed for the frontal ROI (F3, Fz, F4) and the occipital ROI (O1, O2). The electrodes included in the ROIs were selected according to the spatial distribution patterns of e4/non-e4 effects in Sample 1 (Figure 1C).

Figure 2A schematically represents the SEM employed to explore the association between APOE genotypes and the frontal MSE for the EC condition at low, medium, and high time scale AUCs and, correspondingly, for the EO condition (Model 1). The standardized latent MSE variable was regressed onto the dummy variable coding APOE e4 vs. non-e4 carriers. The loadings of the electrode-level factors were fixed to be equal. Table 2 (top) summarizes the results for the frontal ROI. All models had a good fit.

**Table 2.**
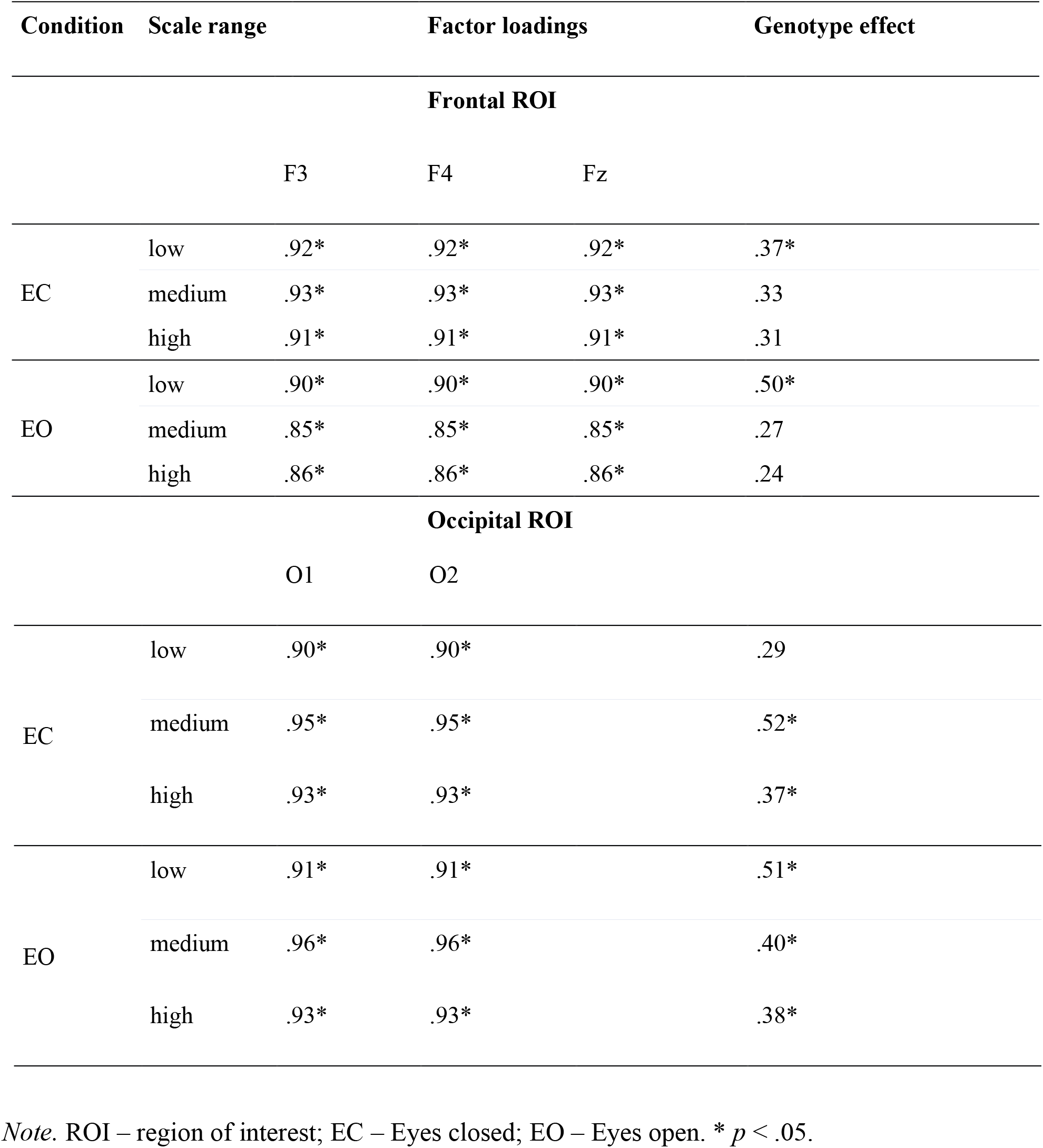
Parameter estimates from Model 1 testing APOE genotype and MSE associations at different ROIs, brain states (EC and EO), and integrated MSE time scales as indicated by AUCs for low, medium, and high time scales.

For the frontal ROI in the EC condition, all loadings of the electrode-specific MSE factors onto the higher-order factor ranged between .8 and .9. The electrodes F3, F4 and Fz were highly consistent with each other in terms of factor loadings. At low, medium and high time scales, the genetic effect was .37 (*p* < .05), .33 (*p* > .05), and .31 (*p* > .05), respectively. Hence, MSE in APOE e4 carriers was larger than in non-carriers at frontal ROI during the EC condition by about one third of a standard deviation. However, the association was only statistically significant for low time scales. Moreover, in accordance with the results provided by Sample 1, the effect size at low scales was slightly larger than at medium and high scales.

In the EO condition at low and medium scales, electrode-specific factor loadings were mostly consistent across all electrode-specific factors. At the low, medium and high scales, the regression weights for genetic associations were .50 (*p* < .01), .27 (*p* > .05) and .24 (*p* >.05), respectively. These associations indicate significantly larger MSE in e4 carriers than in non-carriers at low scales. For medium and high scale AUCs, the genotype effect on MSE failed to reach statistical significance, and the positive effect for high scales was not in line with the discovery Sample 1, which indicates lower MSE in APOE e4 carriers at high scales.

Figure 2B provides a schematic representation of the SEM investigating the APOE genotype effect on MSE in the occipital ROI. All models had a good fit. According to Table 2 (bottom), in the EC condition the regression weights for low, medium and high scales were .29 (*p* > .05), .52 (*p* < .01) and .37 (*p* < .05), indicating higher MSE in e4 carriers as compared with non-carriers. The associations were more pronounced at medium and high scales, which is in line with results from the discovery sample. In the EO condition, there were consistent significant positive genotype associations with MSE at all scales.

Overall, APOE e4 carriers showed higher MSE than non-carriers. The APOE genotype-MSE associations at the frontal ROI were stronger at low scales in both EC and EO conditions. The parietal region showed strong associations in all time scales and in both conditions.

### Associations between APOE e4, MSE and gf

Next, structural equation modelling was conducted to investigate the association between MSE and fluid intelligence, both independently (Model 2), as well as with MSE as mediator (Model 3). Figure 3 displays schematic representations of the models. The latent variable MSE in Figure 3 was indicated in the same way as in Figure 2. The latent variable gf was derived from the three Raven item parcels.

As illustrated in Figure 3A (Model 2), gf was independently predicted by MSE. In Model 2, overall MSE had very weak negative effects on gf at all scales in the EC condition and at the low scale in the EO condition (Table 3). However, the effect was positive at medium and high scales in the EO condition at occipital sites. These results indicate that higher MSE at frontal region and lower scales were weakly associated with lower gf, whereas higher MSE at higher scales and in the EO condition, especially at occipital sites, were weakly associated with higher gf.

**Table 3.** Parameter estimates in the SEMs testing MSE effects on gf (Model 2) and a mediation effect of MSE on the APOE genotype associations with gf (Model 3) for different conditions, time scale ranges and regions of interest.

| Condition | Scale range | Effects in Model 2 | Effects in Model 3 |
| --- | --- | --- | --- |

| Frontal region of interest |  |  |  |  |  |
| --- | --- | --- | --- | --- | --- |
|  |  | gf ~ MSE | MSE ~ APOE | gf ~ APOE | gf ~ MSE |
| EC | low | -.16 | .31 | .49* | -.20 |
|  | medium | -.12 | .31 | .47* | -.15 |
|  | high | -.02 | .27 | .43 | -.04 |
| EO | low | -.09 | .48* | .50* | -.10 |
|  | medium | .00 | .31 | .43 | -.03 |
|  | high | .05 | .19 | .41 | .04 |
| Occipital region of interest |  |  |  |  |  |
|  |  | gf ~ MSE | MSE ~ APOE | gf ~ APOE | gf ~ MSE |
| EC | low | -.22* | .31 | .51* | -.25* |
|  | medium | -.11 | .55* | .49* | -.16 |
|  | high | -.03 | .44* | .44 | -.06 |
| EO | low | -.03 | .47* | .45* | -.07 |
|  | medium | .14 | .51* | .35 | .12 |
|  | high | .23* | .29 | .36 | .21 |

EC and EO = eyes closed and eyes open conditions, respectively; gf = fluid intelligence; MSE = multiscale entropy;

Figure 3B shows Model 3, investigating whether MSE mediates the APOE genotype effect on gf. Model 3 was inspired by our previous work (Li et al., 2019), indicating that APOE e4 carriers with higher education tended to show superior cognitive abilities relative to non-carriers. Given the result that MSE is related at multiple scales with gf, it is of interest to study whether MSE mediates the effect of the APOE genotype on gf. If such a mediation effect exists, the effect of APOE genotype on gf as shown by Li et al. (2019) might be abolished or diminished when MSE is included as a mediator.As indicated on the right side of Table 3, after including MSE as mediator, the direction of the effects of APOE on gf remained unchanged relative to Model 2. Furthermore, the size of the genotype effect on gf did not diminish as compared with the model in which gf was predicted independently by APOE genotype (Li et al., 2019); instead, it slightly increased and reached statistical significance at low and medium scales.

## Discussion

We investigated whether brain signal complexity indicated by MSE differs between young healthy APOE e4 carriers and non-carriers. The results from a discovery sample were confirmed by a replication sample. Furthermore, we assessed how fluid intelligence (gf) is simultaneously related to the APOE genotype and brain signal complexity as a possible mediator between genes and cognitive ability. We argued that even among young healthy individuals, brain signal complexity may be a sensitive neural indicator of APOE genotype differences, which are known to be related to the risk for Alzheimer’s Disease in later life. Therefore, we also tested the association between the APOE genotype and brain signal complexity in relation with fluid intelligence. Our main findings were as follows: 1) The APOE e4 allele was associated with overall higher brain signal complexity across most scales as compared with non-carriers of the e4 allele. 2) The pattern of the association depended on the region of interest on the scalp: At the frontal ROI, the association was strongest at low scales, whereas at the occipital ROI, it was stronger at medium and high scales. 3) Higher fluid intelligence was related with smaller brain signal complexity at low and medium scales, but with larger complexity at high time scales, especially at occipital ROIs in the eyes open condition. These results provide novel insight into the gene-brain-behavior relationships in the context of Alzheimer’s Disease risk.

### Association between brain signal complexity and the APOE e4 risk allele in young adults

For all time scales, APOE e4 carriers were characterized by larger brain signal (EEG) complexity than non-carriers independently of the recording condition (open or closed eyes). At variance with our expectations, the association between APOE e4 genotype and brain signal complexity was not scale-dependent. These results were at variance with an fMRI study by Yang et al. (2014) which failed to identify a difference in brain signal complexity between young APOE e4 carriers and non-carriers. Even though the mechanism through which the APOE genotype affects EEG signal complexity is yet unclear, our results provide evidence about such an effect in young healthy genetic Alzheimer’s Disease risk carriers as compared to non-carriers. Contrary to the commonly accepted knowledge that APOE e4 in older individuals is associated with reduced brain metabolism (Reiman et al., 1996; Small et al., 1995) and functional activity (e.g., Smith et al., 1999), our result in young healthy adults indicate the opposite. This is to say that young risk carriers of Alzheimer’s Disease have more active brains as indicated by brain signal complexity. Future studies taking into account the interaction of genetic Alzheimer’s Disease risk and aging and potential relationships with brain signal complexity might reveal further insights into the gene-brain-behavior associations in the context of Alzheimer’s Disease risk.

In our study, APOE e4 association with brain signal complexity was studied with both a discovery and a replication sample. The result was mostly consistent across the two samples at low scales, frontal ROI under EC condition, as well as medium and high scales, occipital ROI under both EC and EO conditions. However, the association was not entirely consistent across the discovery and replication samples. For example, the association of MSE at low scales and frontal ROI in in the eyes open condition was not present in the discovery sample but was significantly positive in the replication sample. The association of MSE at high scales in the frontal ROI was mostly negative in the discovery sample but positive and much smaller in effect size in the replication sample. This discrepancy might be attributed to the small size of the discovery sample and underscores the importance of replication trials. Also, even though the discovery and replication samples were separately recruited from population with similar demographic factors (Huffman et al., 2018), the sample collection and EEG recording procedures were designed for different study aims, while the EEG lab parameters were not unified.

In general, the present results suggest that occipital-posterior brain signal complexity is most robustly associated with APOE genotype differences in young adulthood. Indeed, in previous fMRI studies (e.g. Dennis et al., 2009; Coughlana et al., 2020; Yang et al., 2014), the posterior cingulate was frequently reported to be more activated in APOE e4 carriers as compared with non-carriers.

### Association between brain signal complexity and fluid intelligence in young adults

The present results suggest that the sign of the association between brain signal complexity and fluid intelligence varies across scale level, being negative at low scales and positive at high scales. According to this result, the low and high scale MSE estimations are related oppositely. According to the interpretation by Costa et al. (2005), Multiscale Entropy estimated at lower time scales capture regularity/randomness in high frequency fluctuations. The coarse graining procedure is intended to filter out high-frequency random fluctuations; hence, higher scale entropy reflects temporally long-range correlations, which characterize the complexity at low frequency of time series structure. Additionally, coarse graining will not affect the signal components for which entropy would potentially characterize real complexity (Costa et al., 2005). In other words, the interpretation of multiscale entropy associations with brain function should be based on the combination of low and high time scales characteristics. This reasoning is supported by multiple studies. For example, an fMRI study revealed that multiscale entropy and functional connectivity are negatively correlated at low scales and positively correlated at high scales (McDonough and Nashiro, 2014; Liu et al., 2019). According to their studies (McDonough and Nashiro, 2014; Liu et al. 2019), MSE captures different scale properties, the low and high scale variations are related through functional connectivity, indicating the broad spectral properties of EEG. This may provide an explanation for our results, that fluid intelligence accounted by functional connectivity was associated with low and high scale MSE in opposite ways.

### Brain signal complexity and the APOE e4 association with cognition

In our previous study (Li et al., 2019) which relied on purely cognitive measures from the present replication sample, fluid intelligence was regressed onto the APOE genotype. Results from the same study sample indicated higher gf in e4 carriers as compared with non-e4 carriers but the difference was not significant (the effect estimate was .23, *p* = .17). Here, the APOE e4 effect on gf increased and reached significance after individual differences in brain signal complexity were controlled for. This result suggests that brain signal complexity does not mediate the relationship between APOE e4 and fluid intelligence. In other words, APOE is not associated with gf via brain signal complexity and the association between APOE e4 and gf exists also within groups of similar brain signal complexity levels. Therefore, other underlying neural process than brain signal complexity may account for the pathway between genes and fluid intelligence which needs to be identified in future research.

Brain signal complexity may have acted as a suppressor variable (Conger, 1974), as the effect of the other predictor, APOE genotype, on the dependent variable, fluid intelligence, increased after brain signal complexity was included in the regression model. The suppressor effect seems to be stronger at low scale, because the increase of the relationship between APOE and gf was stronger when low scale MSE was controlled for, as compared with the case when high scale MSE was accounted for. We explain this suppressing effect of low scale MSE on fluid intelligence as that higher MSE at low scale may suggest stronger local processing (Vakorin et al., 2011), which may be less relevant for fluid intelligence which requires more integrated processing (Gard et al., 2014).

One limitation of the current study is that we did not take education level into consideration. In our previous study (Li et al., 2019), we investigated the interaction between the APOE genotype and education level on cognitive performance and detected negative APOE e4 effect on cognitive performance among individuals with lower education level and weak positive APOE e4 effect on cognitive performance among persons with higher education. The current study was applied to a selective sample with higher education as compared with the sample described in Li et al. (2019). Thus, the detected gene – brain – behavior relationship and related conclusions can only be drawn for a high education population. Future studies are needed to take the complex effect of environmental factors, such as education into account.

### Conclusions

We found significantly positive APOE e4 associations with brain signal complexity at frontal and occipital brain regions among young healthy adults, and the signal complexity may slightly be associated with individual differences in fluid intelligence. However, brain signal complexity does not mediate the APOE genotype effect on fluid intelligence. These results may contribute to understanding the neural process underlying the APOE effect on behavior.

## Supporting information

MSE data

## Supplementary material

It is agreed that MSE calculation usually needs very long time series for high reliability, because as scale factor increases, the estimation variance of sample entropy will grow quickly since the data series is highly coarse-grained. Here we provide a systematic assessment of the relation between reliability of scale-wise MSE measurements and length of the EEG segments used for MSE calculation. The MSE reliability estimates were calculated as Composite Reliability (Raykov, 1997).

**Figure S1.**
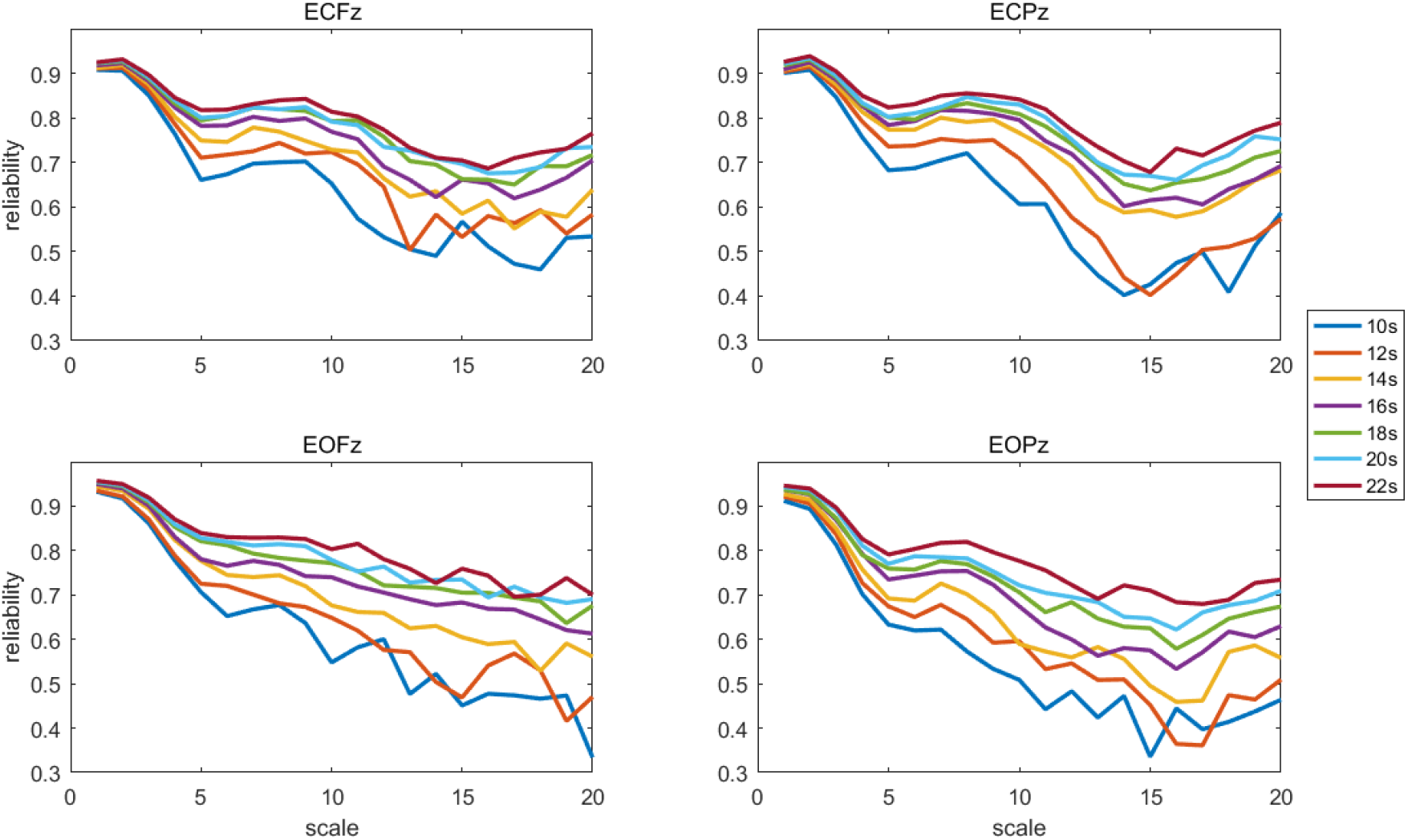
Reliabilities for scales 1-20 for MSE measurements estimated for different lengths of EEG data. ECFz: Fz channel, eyes closed condition. ECPz: Pz channel, eyes closed condition. EOFz: Fz channel, eyes open condition. EOPz: Pz channel, eyes open condition.

